# OGUs enable effective, phylogeny-aware analysis of even shallow metagenome community structures

**DOI:** 10.1101/2021.04.04.438427

**Authors:** Qiyun Zhu, Shi Huang, Antonio Gonzalez, Imran McGrath, Daniel McDonald, Niina Haiminen, George Armstrong, Yoshiki Vázquez-Baeza, Julian Yu, Justin Kuczynski, Gregory D. Sepich-Poore, Austin D. Swafford, Promi Das, Justin P. Shaffer, Franck Lejzerowicz, Pedro Belda-Ferre, Aki S. Havulinna, Guillaume Méric, Teemu Niiranen, Leo Lahti, Veikko Salomaa, Ho-Cheol Kim, Mohit Jain, Michael Inouye, Jack A. Gilbert, Rob Knight

## Abstract

We introduce Operational Genomic Unit (OGU), a metagenome analysis strategy that directly exploits sequence alignment hits to individual reference genomes as the minimum unit for assessing the diversity of microbial communities and their relevance to environmental factors. This approach is independent from taxonomic classification, granting the possibility of maximal resolution of community composition, and organizes features into an accurate hierarchy using a phylogenomic tree. The outputs are suitable for contemporary analytical protocols for community ecology, differential abundance and supervised learning while supporting phylogenetic methods, such as UniFrac and phylofactorization, that are seldomly applied to shotgun metagenomics despite being prevalent in 16S rRNA gene amplicon studies. As demonstrated in one synthetic and two real-world case studies, the OGU method produces biologically meaningful patterns from microbiome datasets. Such patterns further remain detectable at very low metagenomic sequencing depths. Compared with taxonomic unit-based analyses implemented in currently adopted metagenomics tools, and the analysis of 16S rRNA gene amplicon sequence variants, this method shows superiority in informing biologically relevant insights, including stronger correlation with body environment and host sex on the Human Microbiome Project dataset, and more accurate prediction of human age by the gut microbiomes in the Finnish population. We provide Woltka, a bioinformatics tool to implement this method, with full integration with the QIIME 2 package and the Qiita web platform, to facilitate OGU adoption in future metagenomics studies.

**Importance:** Shotgun metagenomics is a powerful, yet computationally challenging, technique compared to 16S rRNA gene amplicon sequencing for decoding the composition and structure of microbial communities. However, current analyses of metagenomic data are primarily based on taxonomic classification, which is limited in feature resolution compared to 16S rRNA amplicon sequence variant analysis. To solve these challenges, we introduce Operational Genomic Units (OGUs), which are the individual reference genomes derived from sequence alignment results, without further assigning them taxonomy. The OGU method advances current read-based metagenomics in two dimensions: (i) providing maximal resolution of community composition while (ii) permitting use of phylogeny-aware tools. Our analysis of real-world datasets shows several advantages over currently adopted metagenomic analysis methods and the finest-grained 16S rRNA analysis methods in predicting biological traits. We thus propose the adoption of OGU as standard practice in metagenomic studies.

## Introduction

The rapidly developing field of shotgun metagenomics has inherited many analytical tools from the more mature field of 16S rRNA gene amplicon studies. For example, diversity analyses provided in platforms such as QIIME 2 (1) can be used for metagenomic analyses. To date, the typical metagenomics workflow starts with taxonomic profiling, which estimates the taxonomic composition of microbial communities by matching sequencing data against a reference database (2). The resulting matches are compiled into an unstructured feature table, with values usually in the form of relative abundances of taxonomic units at a fixed rank (e.g. genus or species level), followed by relevant statistical analyses.

In contrast, the current standard for 16S rRNA analysis involves more advanced feature extraction, including construction of amplicon sequence variants (ASVs), which have replaced operational taxonomic units (OTUs) to deliver the finest-possible resolution from amplicon data (3). Phylogeny-aware algorithms such as UniFrac (4) have been widely-adopted to model community diversity while considering how features interrelate owing to the accessibility of reference phylogenies (5, 6), and the availability of *de novo* and *a priori* phylogenetic inference methods (7). This wisdom should be adopted as well to metagenomics. Thanks to the advances in efficient sequence alignment algorithms, and the expansions of reference genome databases (8, 9) and phylogenomic trees (10, 11), it is now possible and increasingly preferable to develop a fine-resolution, structured data analysis strategy in shotgun metagenomics.

Therefore, we propose an alternative method for constructing metagenomic feature tables, in which features are no longer taxonomic units, but individual reference genomes from a database, and the feature counts are the number of sequences aligned to these genomes. We refer to such features as Operational Genomic Units (OGUs). This term, in an echo of OTU but replacing “taxonomic” with “genomic”, highlights the nature of the genome-based, taxonomy-free analysis. Meanwhile, “operational” indicates that this method does not rely on the direct observation of member genomes of the community, but uses pre-defined reference genomes as a proxy to model the community composition. However, like ASVs, OGUs are exact and do not rely on similarity thresholds as OTUs do.

An OGU table represents the finest-grained resolution of observed genomes in a microbial community relative to the reference database. As such it can be used to quantify the community structure and relationships in correlation with biological traits. It can also work well with cost-efficient “shallow” shotgun metagenomics (12), where limited sequencing depth (even below the previously recommended lower threshold of 500,000 sequences per sample) is adequate for assessing community structure. It further empowers tree-based analyses, such as UniFrac and phylofactorization, which is enhanced by using the “Web of Life” (WoL) reference phylogenetic tree that we recently developed to describe accurate evolutionary relationships among genomes (10).

We have implemented the method for generating OGU tables in the open-source bioinformatics tool, Woltka (https://github.com/qiyunzhu/woltka). This program serves as a versatile interface connecting choices of upstream sequence aligners (such as Bowtie2 and BLAST) and downstream microbiome analysis pipelines (such as QIIME 2). In addition to the standalone program, the package ships with a QIIME 2 (1) plugin to facilitate adoption and integration into existing protocols. We have also made this method available through the Qiita web analysis platform (13) as part of the standard operating procedure for shotgun metagenomic data analysis, thereby enabling massive reprocessing and subsequent meta-analysis of metagenome datasets with OGUs. Thus far, we have applied the OGU method to re-analyze all public and private metagenomic datasets hosted on Qiita, totaling 143 studies and 57,063 samples, as of Mar 3, 2021.

Our team and collaborators have applied prototypes of the OGU method in multiple microbiome and multiomics studies and have obtained biologically relevant results (e.g., (14–16)). In this article, we systematically introduce the principles and practices of the OGU method, demonstrate its efficacy in one synthetic and two real-world microbiome datasets, and compare it with state-of-the-art metagenome analysis approaches and the alternative data type (16S rRNA gene amplicons). Given our findings, we propose the adoption of OGUs as a good practice in metagenomic analyses.

## Results

### OGUs maximize resolution of community structures

The rationale and benefits of the OGU method are demonstrated with a synthetic case study illustrated in Fig. 1, with the underlying feature tables provided in Table S1. In this simple case, three metagenomes with 12 sequences each were aligned to 10 reference genomes, which were hierarchically organized by taxonomy (left) or by phylogeny (right) (Fig. 1A). Beta diversity was calculated on feature tables at different levels: either on taxonomic units at the rank of genus or species, or directly on reference genomes (i.e., OGUs) without the need for giving them taxonomic labels.

**Figure 1.**
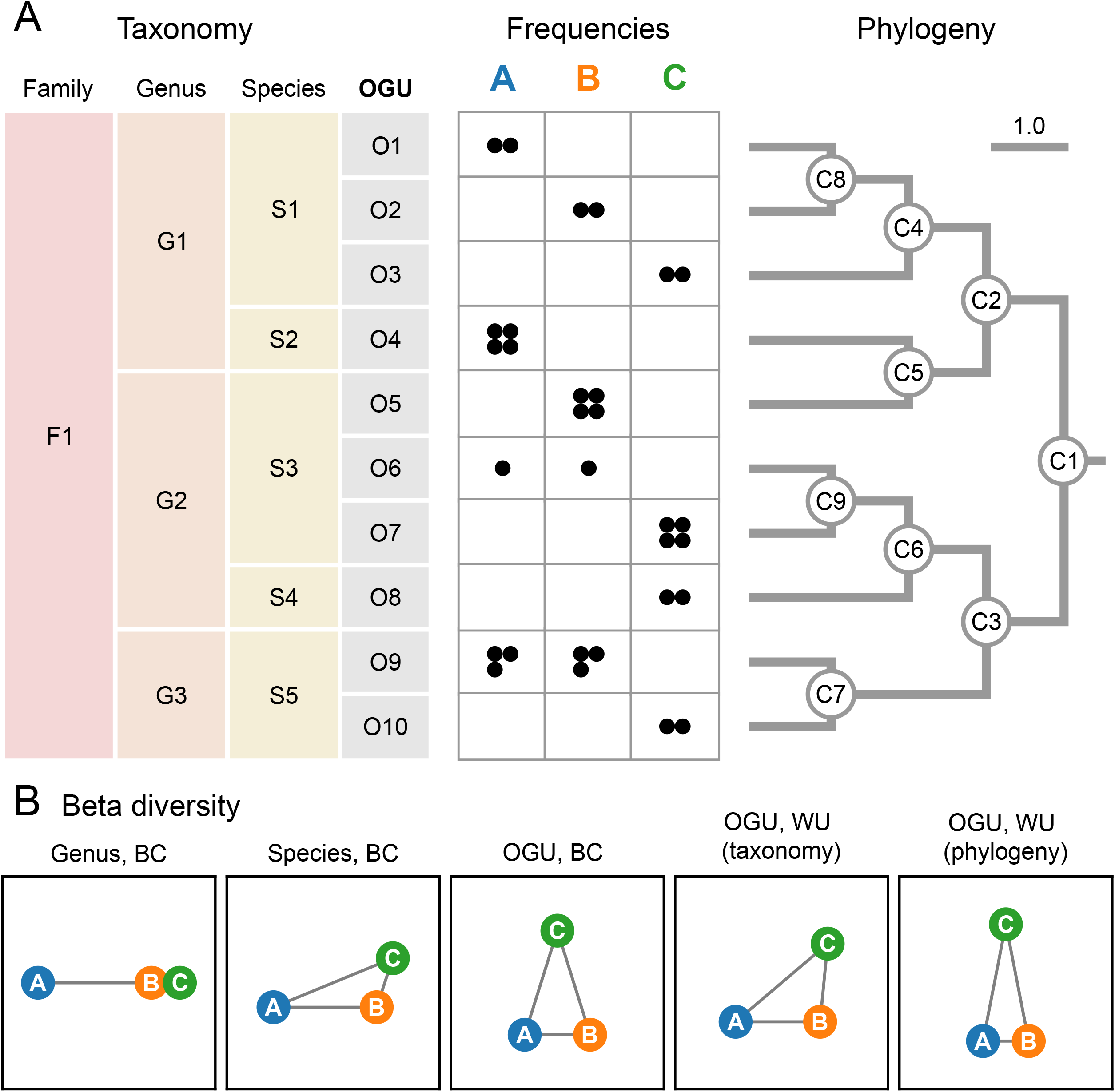
Feature resolution impacts community structure analysis even in small conceptual examples. **A**. A synthetic dataset involving three microbial communities, each of which having 12 unique read hits, as represented by black circles in the frequency table, to a total of 10 reference genomes (OGUs), classified under five species, three genera and one family, as noted to the left. A phylogenetic tree of the 10 genomes is shown on the right. In this simplified case, the phylogeny is not much more complex than the taxonomy (with three more edges); however, the taxonomic assignment and the phylogenetic placement of genome O5 are not consistent. **B**. Beta diversity of the dataset. The three samples (circles) are connected by edges representing the pairwise distances calculated by Bray-Curtis (BC) or weighted UniFrac (WU) on the frequency table. For the latter measure, either the taxonomy or the phylogeny was used to quantify the hierarchical relationships among OGUs, as noted in the parentheses. The edge lengths were normalized so that their sum is equal in each graph. This synthetic case study demonstrates that different resolutions of features and feature structures can lead to very different conclusions regarding sample relationships.

As demonstrated (Fig. 1B), the genus-level analysis, which had the lowest resolution (three genera), yielded spurious proximity between samples B and C, as relative to sample A, largely determined by the differential abundance of genera G1 and G2. The species-level analysis with moderately higher resolution (five species) was able to bring A closer to B and C, mainly contributed by the identical frequencies of species S1, which could not be revealed at the genus level. The OGU-level analysis, having the highest resolution (10 features), revealed the separation between B and C due to distinct OGU composition, despite similar species counts (e.g., O5 and O7 have different counts within S3), and the proximity between A and B due to shared OGUs (O6 and O9). Additional structure was revealed by using the UniFrac metric, which considers the hierarchical relationships among features, hence further joining samples (here A and B) sharing longer branches in the phylogenetic tree (even by different OGUs, such as O1 and O2) and separating those sharing shorter ones. Taxonomy may serve as a replacement of phylogeny, but it has a lower resolution than phylogeny (e.g., O1 and O2 are evolutionarily closer to each other relative to O3 but taxonomy cannot reveal this), and sometimes does not reflect the true evolutionary relationships among organisms (e.g., O4 and O5 are here placed in different genera), which can impact the accurate modeling of community structures.

In summary, this example illustrates the need for increasing resolution in order to better understand the diversity of microbial communities. This “resolution” has two dimensions of meaning: first, the quantity of features representing individual microbiomes; second, the granularity and accuracy of the hierarchy— if any—that defines the relationships among individual features.

### OGUs accurately represent body environment and host sex associated microbiome patterns

We demonstrated the typical use of the OGU method on the classic Human Microbiome Project (HMP) shotgun metagenomic dataset (17), which contains 210 metagenomes sampled from seven body sites of male and female human subjects. We subsampled each metagenome to one million paired-end reads—a sampling depth close to the recommended lower threshold (500k reads) for “shallow” shotgun sequencing (12). The sequences were aligned to the WoL reference genome database (totaling 10,575 bacterial and archaeal genomes) and the alignments were processed using Woltka, resulting in an OGU table with 6,220 features (reference genomes) (Fig. S1A). Beta diversity analysis using the weighted UniFrac metric with the WoL reference phylogeny was performed on the OGU table (Fig. 2). For comparison, we analyzed the dataset using the currently adopted method (CAM) (e.g., (17)): using Bray-Curtis on a species-level taxonomic profile. We exemplified the CAM by using the profile inferred by Bracken (18) on the same WoL database (Fig. 2), but also tested and reported the results of SHOGUN (19), Centrifuge (20), and MetaPhlAn (21) (Fig. S1).

**Figure 2.**
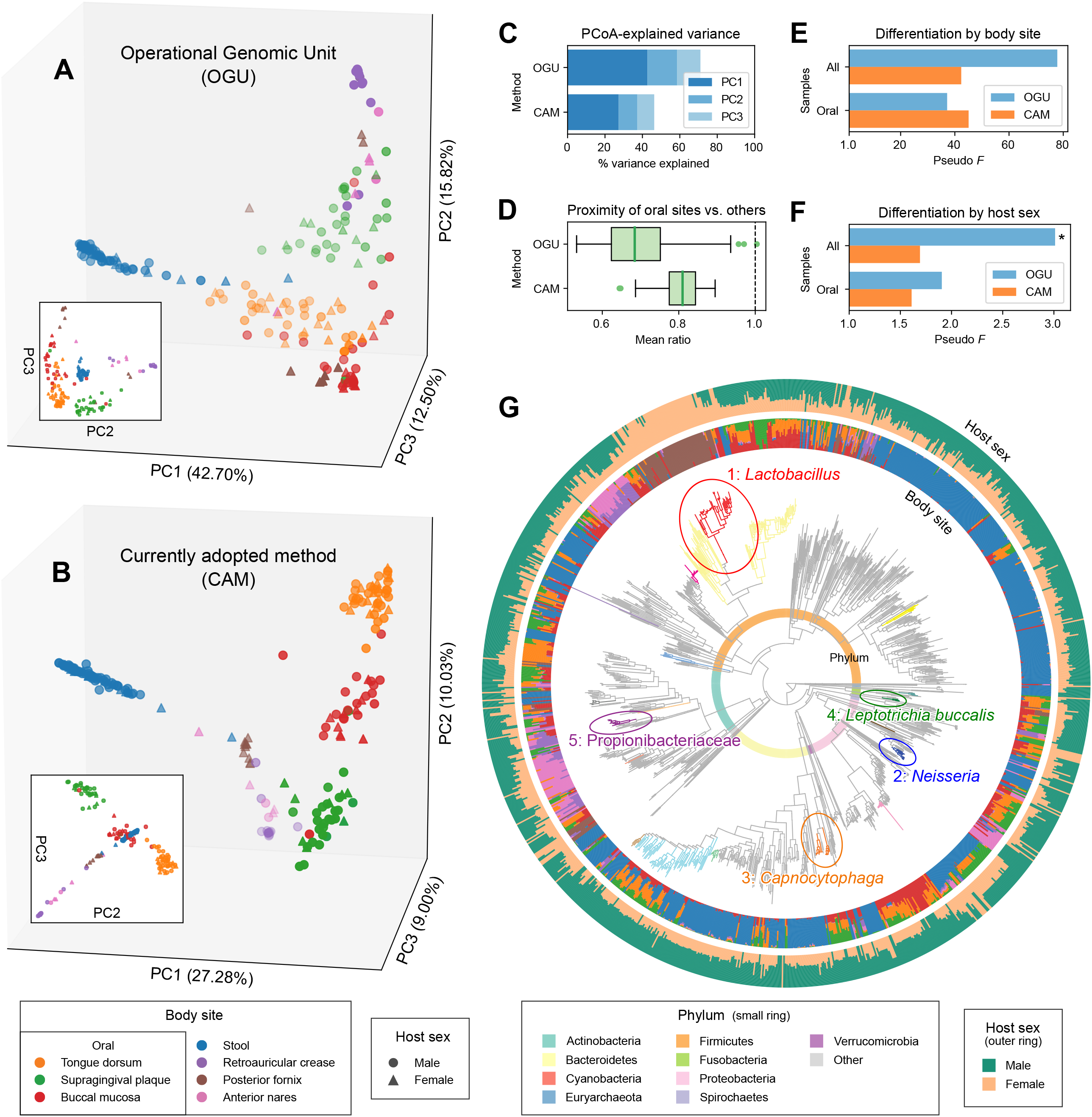
Analysis of the HMP metagenomes reveals clustering by body environment and differentiation by host sex. Beta diversity analysis was performed on 210 samples subsampled to one million paired-end shotgun reads each. **A**. PCoA by the method proposed in this study (OGU): weighted UniFrac metric calculated with the WoL reference phylogeny based on the OGU table. Samples (dots) are colored by body site and shaped by host sex. **B**. PCoA using the current adopted method (CAM): Bray-Curtis calculated on species-level taxonomic units identified by Bracken, which shows a diagonal pattern that aligns all samples of the four non-oral body sites in one plane (also see Figs. S2B and S3). **C**. Proportions of community structure variance explained by the first three axes of PCoA. **D**. Mean ratio of the beta diversity distances from any oral sample to a sample of the two other oral sites versus to that of non-oral body sites. The lower the mean ratio is, the more similar communities of the three oral sites are to each other in the background of multiple body environments. The bold line in each box represents the median. The whiskers represent 1.5 IQR. **E** and **F**. PERMANOVA pseudo-*F* statistics indicating the differentiation of community structures by body site (**E**) and by host sex (**F**). The larger *F* is, the more distinct the community structures are between groups versus within groups. The *y*-axis is aligned to *F*=1.0 which indicates no difference. For **E**, all statistics have a *p*-value of 0.001. For **F**, an asterisk (*) indicates *p*-value ≤ 0.05. **G**. Differentially abundant phylogenetic clades by host sex inferred using PhyloFactor and visualized using EMPress on the WoL reference phylogeny. The tree was subsetted to only include OGUs detected in the dataset. The top 20 clades by effect size are colored (full details provided in Figs. S4-5). The top five clades are numbered 1 through 5 by decreasing effect size, circled, and labeled with corresponding taxonomic annotations. The small color ring represents phylum-level annotations. The inner and outer barplot rings indicate the OGU counts split by body site (using the same color scheme as in A and B) and by host sex, respectively.

Principal Coordinates Analysis (PCoA) of OGUs (Figs. 2A and S2A), with the first three axes explaining 71.01% of community structure variance (Figs. 2C and S1B), revealed that microbiomes were clustered mainly by the body site from which they were sampled, which overshadowed clustering by host sex, if any. This pattern is largely consistent with the previous report (17). The PCoA plot by CAM (Figs. 2B and S2B, also see S3), although with less explained variance (46.30%) (Figs. 2C and S1B), also displayed a clustering-by-site pattern. However, it is notable from the plot that sample clusters are aligned diagonally—a typical pattern indicating the saturation of distances caused by the inadequacy of shared features (species) among body sites (22) (Figs. 2B and S2B). This characteristic limits the power of resolving community diversity.

Permutational multivariate analysis of variance (PERMANOVA) of the beta diversity distance matrices suggested that all methods were able to clearly differentiate samples by body site (*p*=0.001), with OGU generating the strongest statistic (Figs. 2E and S1C) (OGU: *F*=77.82; CAM: *F*=42.36). The distinction by host sex was less obvious. Only OGU was able to distinguish microbiome by sex (*F*=3.011, *p*=0.013), whereas CAM failed to distinguish sex with statistical significance (*F*=1.692, *p*=0.086) (Figs. 2F and S1E-F). This demonstrated the power of the OGU method in capturing subtle but relevant trends, even when another primary factor (body site) is driving most of the community diversity. Three of the seven body sites are located in the oral environment: tongue, teeth and buccal mucosa (Fig. 1A, B). They together indicate weaker differentiation by sex (OGU: *F*=1.905, *p*=0.099; CAM: *F*=1.610, *p*=0.130) (Figs. 2F and S1G-H). In parallel, we reason that sites sharing the same environment likely have higher microbial connections. To test this effect, we calculated the relative distance between the three oral sites versus oral sites to non-oral sites. This distance is significantly smaller with OGU (0.699 ± 0.098, mean and std. dev., same below) than with CAM (0.808 ± 0.051) (two-tailed paired *t*=-14.398, *p*=2.57e-26) (Figs. 2D and S1D), suggesting that OGU is more effective at relating subgroups of samples with shared properties.

The OGU table plus the WoL tree further enabled differential abundance analysis using the phylogenetic factorization method (23) (Figs. S4-5). The result was visualized and analyzed using the recently released massive tree visualizer EMPress (24) (Fig. 2G). It revealed that the phylogenetic clade separated by Factor 1 represents the genus *Lactobacillus*, contained in predominantly posterior fornix samples from female hosts, which is expected (25). Meanwhile, Factor 2 (genus *Neisseria*), Factor 3 (genus *Capnocytophaga*) and Factor 4 (species *Leptotrichia buccalis*) are more frequently observed in the oral sites of male hosts. For comparison, we applied the tree-free method ANCOM (26) on the taxonomic profiles generated by alternative methods (Table S2). At genus level, all four methods were able to capture only *Lactobacillus*, consistent with our Factor 1. However, at species and OGU levels, results were discordant between methods and no method reported any *Lactobacillus* sp., again showing the limitations of confining analyses to taxonomic ranks without phylogenetic information.

Finally, we assessed the efficacy of OGUs along a gradient of decreasing sampling depths. The correlation between the original OGU table (from one million paired-end reads) and each of the subsampled OGU tables was consistently high. A Pearson’s *r* of 0.961 ± 0.0726 (mean and std. dev., same below) was retained even at the sampling depth of 200 (Fig. S6A). The PCoA clustering pattern largely remained the same at all sampling depths (Fig. S7). The oral-vs-other relative distance (see above) retained a Pearson’s *r* of 0.971 ± 0.00613 when sampling depth was 200 (Fig. S6B). The PERMANOVA *F*-statistics calculated based on 10 replicates of random subsampling were close to the original statistic and largely stable down to very low sampling depths. The mean difference from the original statistic was still within 5% at the sampling depth of 1,000 for body site (3.349 ± 1.361, unit: percentage of the original statistic, same below), or 500 for host sex (2.680 ± 5.473) (Fig. S6C-D). These findings suggest that the OGU method remains valid even on very shallow metagenomic samples, including those that would otherwise be considered unusable for typical metagenomic analyses.

### OGUs improve prediction of host age from the gut microbiome

We next analyzed 6,430 stool samples collected through a random sampling of the Finnish population using both 16S rRNA gene amplicon sequencing and shallow shotgun metagenomic sequencing. This “FINRISK” study (27) provides an opportunity to explore the dependency of feature sets (e.g. taxonomic levels and data source: 16S rRNA amplicon vs. shotgun metagenomic data) on the prediction accuracy of a machine learning model on the targeted phenotype (e.g., age). We quantitatively examined the impact of taxonomic level of microbiome features on the empirical error (mean absolute error, or MAE) in predicting human chronological age using a Random Forests regressor (28), constructed using 5-fold cross-validation.

Our results (Fig. 3A) showed the prediction accuracy continued to improve, resulting in lower absolute errors with finer microbial feature classification levels. Shotgun data outperformed 16S data at all levels, and was able to reduce MAE to less than 10 years at the genus level or below. At the lower limit of both 16S and shotgun data, we achieved an MAE of 9.581 ± 0.116 years (mean and std. dev., same below) with OGUs (Fig. 3B), whereas ASVs, the highest possible resolution allowed by 16S data, resulted in a higher MAE of 10.110 ± 0.103 years (two-tailed *t*=-7.25, *p*=8.81e-5). Meanwhile, using the species-level profile inferred by Bracken, we also obtained a higher MAE of 10.273 ± 0.089 years (vs. OGU: two-tailed *t*=-10.59, *p*=5.53e-6) (Fig. S8). Decreasing sequencing depth did not reduce the age prediction accuracy for individual samples (Fig. S9). For example, samples with 320-366k metagenomic sequences (2nd bin from low end in the figure) had an MAE of 9.290 ± 6.378 years, whereas samples with 1,386-1,931k sequences (2nd bin from high end) had an MAE of 10.118 ± 6.086 years, which were not significantly different (two-tailed *t*=-1.37, *p*=0.170). We then explored which OGUs contributed to the superior performance in age prediction as compared to 16S rRNA ASVs. Therefore, we identified a reduced set (*n*=128) of the most important OGUs that can maximize the prediction accuracy via a recursive feature elimination approach (Fig. S10). Among these important features, a few gut microbial strains increased in abundance with aging, such as multiple strains from *Streptococcus mutans, Eubacterium sp*. (Figs. 3C, S11-12). Remarkably, those *Streptococcus* spp. are typically located in the oral cavity yet can be over-represented in the gut of elderly individuals, suggesting potential microbial transmissions between oral and gut microbiomes related to typical aging in a large population (29, 30). Next, we also identified a few microbial OGUs that were under-represented in the elderly, such as *Anaerostipes hadrus* DSM 3319 and members of *Bifidobacterium*, including *B. longum* NCC2705 and *B. saguini* DSM 23967 Bifsag. Many of these important taxonomic features were not identified in the 16S data, putatively because the partial sequences of a 16S rRNA gene cannot provide sufficient resolution to distinguish species or strains. For example, a few 16S rRNA ASVs annotated with *Lachnospiraceae* have been associated with aging and were identified in either this or past studies (31), whereas our method identified several OGUs (*Anaerostipes hadrus* DSM 3319) within the family of *Lachnospiraceae* that exhibited strong predictive powers for discriminating aging.

**Figure 3.**
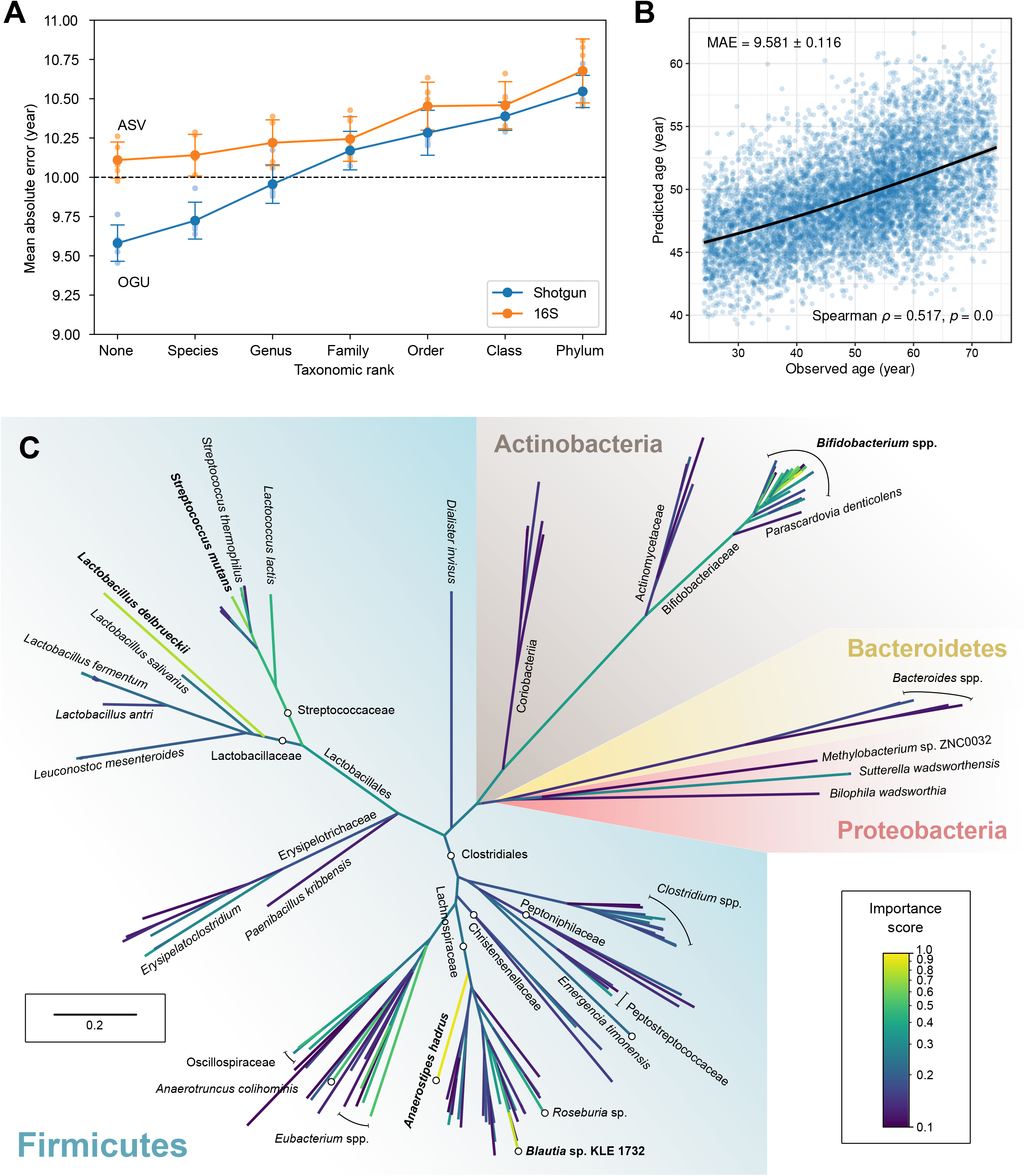
Analysis of the FINRISK metagenomes showing superior prediction accuracy over taxonomic units and 16S rRNA data. **A**. The empirical error (mean absolute error, MAE) in predicting host chronological age using microbiome features at distinct taxonomic ranks in paired 16S rRNA amplicon and shotgun metagenomics data with a Random Forests regressor. “None” represents the taxonomy-free, finest-possible level (ASV for 16S, OGU for shotgun). Small circles indicate MAEs in all iterations of five-fold cross validation. Large circles and error bars indicate the mean and standard deviations of the five MAEs. **B**. Scatter plot of the actual age vs. the predicted age by the best-performing model with OGU features in the five-fold cross-validation. The black line was generated using ggplot2’s local polynomial regression fitting. **C**. Phylogenomic tree of 169 OGUs with importance score ≥ 0.1 in the prediction model. The tree was subsampled based on the WoL reference phylogeny, and drawn to scale (branch lengths represent mutations per site). Branch colors indicate the mean importance score of all descendants of the clade. Taxonomic labels are displayed where needed. Circles and lines with stops are displayed where needed to assist location of taxonomic labels to target branches or clades.

## Discussion

The OGU method introduced in this article provides a way to maximize the resolution of feature tables by directly considering reference genomes without the reliance on taxonomic classification in shotgun metagenomics studies. Although the strategy of taxonomy-free community structure analysis has been widely adopted in 16S data analysis (e.g., ASV or *de novo* OTU clustering), it remains underexplored in metagenomics, largely due to the difficulties in defining and quantifying “features” without using an *a priori* classification system. Our study shows that sequence alignment hits to individual reference genomes can be used as the minimum unit for features, referred to as OGUs.

Through comparative analysis of OGU and alternative methods using a synthetic case study and two real-world microbiome studies, we demonstrated that classical high-dimensional statistics and machine learning methods developed and matured in the field of 16S rRNA gene amplicon analysis can be directly applied to OGUs to provide biologically relevant insights. The OGU results often are superior to currently adopted metagenomic classification methods and ASV analysis of the 16S rRNA data.

Meanwhile, we showed that the use of taxonomic units as features, as many researchers have been practicing to date, has conceptual and performance limitations compared with the OGU method, particularly at higher taxonomic ranks due to the loss of resolution.

The independence from taxonomy further enables the utilization of explicit phylogenetic trees. A researcher can choose from pre-computed reference phylogenies, such as the one we introduced in the “Web of Life” (WoL) project (10), or custom phylogenomic trees computed from *de novo* construction or placement, through tools such as PhyloPhlAn3 (32) and DEPP (33), which are scalable to large numbers of genomes. This connects evolutionary biologists’ efforts in updating the tree of life (e.g., (10, 11, 34)), computational biologists’ efforts in forging phylogeny-aware methods (e.g., UniFrac and PhyloFactor), and microbiome scientists’ pursuits of relating high-dimensional microbiome data with biology.

Taxonomy, despite being relatively coarse-grained and error-prone as a classification system, may serve as an implicit replacement of phylogeny if the latter is not available. We tested this idea by applying UniFrac to an artificial taxonomic tree with constant branch lengths between ranks (analogous to (35)). Although this treatment is controversial, because taxonomic ranks do not directly indicate evolutionary distances, we did observe improvement compared to not using a tree (Fig. S13). Although there have been remarkable efforts for curating taxonomy using phylogenetics, however, the number of taxonomic ranks is limited (typically 7 to 8), and can constrain the topology for an ever-growing number of sequenced genomes. For example, the current release (R95) of GTDB (36) has 31,910 species clusters, constituting a taxonomy tree of 45,502 vertices, whereas NCBI RefSeq and GenBank host 977,729 unique genomes as of March 30, 2021, and a fully resolved phylogenetic tree of them can theoretically have 1,955,456 vertices. The history of 16S rRNA studies (7) is repeating itself in whole-genome studies, such that building a phylogeny is not only advantageous but often more feasible than defining taxonomy, and the OGU method powerfully provides an analogous extension to shotgun sequencing studies. As a new notion to microbiome research, OGU’s properties in statistical analyses has yet to be characterized in a large number of studies, as was done for 16S rRNA ASVs. Unique challenges in shotgun metagenomics may impact analyses that were designed for 16S rRNA data. For example, very-low-abundance false positive assignments, which are prevalent from typical metagenomic classifiers, may impair the accuracy of the recovered community composition (37). A typical treatment is to only consider features with relative abundance above a given threshold in each sample (37). While we provide this function in Woltka to facilitate user’s preferences, our tests suggested that the result of an OGU analysis is highly stable against a wide range of filtering thresholds when using abundance-based metrics (weighted UniFrac and Bray-Curtis), as compared with presence/absence-based metrics (unweighted UniFrac and Jaccard) (Fig. S14). This observation implies the OGU method is robust to noise commonly introduced into metagenomic datasets from many low abundance observations.

The robustness of an OGU analysis is only limited by the comprehensiveness of the reference. Despite that available genomic data have grown to an enormous volume, the size of a reference genome database that can be realistically used in a metagenomic analysis with typical computing facilities is circumscribed, limiting the increase of resolution beyond sub-species levels. Balancing alignment accuracy and database content is therefore an important consideration in designing the analytical strategy. The algorithm we previously designed and used in the WoL database to maximize the covered biodiversity given a fixed number of genomes (10) may be beneficial in this situation, but its efficacy needs to be further tested in the background of various biospecimens and biological questions.

Leaderboard sequencing may also be a useful strategy for iteratively augmenting the reference database with the common genomes in each sample (38). In the long run, efforts to improve algorithms, increase database coverage, and improve computing efficiency are all needed to facilitate effective advances in the field of metagenomics, and the OGU method provides an important step forward in that direction.

## Materials and Methods

### Protocol details

The OGU method is flexible to the type of sequence alignment. The recommended protocol, which is also the protocol demonstrated and benchmarked in this article, is as follows: Shotgun metagenomic sequencing data were aligned against the WoL reference genome database using SHOGUN v1.0.8 (19), with Bowtie2 v2.4.1 (39) as the backend. This process is equivalent to a Bowtie2 run with the following parameters:

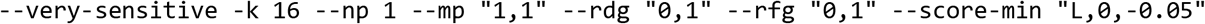

The sequence alignment is treated as a mapping from queries (sequencing data) to subjects (reference genomes). It is possible that one sequence is mapped to multiple genomes (up to 16 using the aforementioned Bowtie2 command). In this scenario, each genome is counted 1 / *k* times (*k* is the number of genomes to which this sequence is mapped. The frequencies of individual genomes were summed after the entire alignment was processed, and rounded to the nearest even integer. Therefore, the sum of OGU frequencies per sample is nearly (considering rounding) equal to the number of aligned sequences in the dataset. The output feature table has columns as sample IDs, rows as feature IDs (OGUs), and cell values as the frequency of each OGU in each sample. This table is ready to be analyzed using software packages such as QIIME 2 (1).

### Implementation

The OGU method is implemented in the bioinformatics tool Woltka (Web of Life Toolkit App), under the BSD-3-Clause open-source license. The program is written in Python 3, following high-quality software engineering standards. Its unit test coverage is 100%. The source code is hosted in the GitHub repository: https://github.com/qiyunzhu/woltka, together with instructions, tutorials, command-line references, and test datasets. The program has been included in the Python Package Index (PyPI). In addition to the standalone Woltka program, a QIIME 2 (1) plugin is included in the software package.

Woltka automatically recognizes and parses multiplexed or per-sample sequence alignment files, either original or compressed using Gzip, Bzip2 or LZMA algorithms. It supports three alignment file formats: 1) SAM (Sequence Alignment Map) (40), which is supported by multiple short read alignment programs, such as Bowtie2 (39), BWA (41) and Minimap2 (42); 2) the standard BLAST (43) tabular output format (“-outfmt 6”), which is supported by multiple sequence alignment programs, such as BLAST, VSEARCH (44) and DIAMOND (45); 3) A plain mapping of query sequences to subject genomes, which is customizable to adopt other tools and pipelines.

In addition to OGU table generation, Woltka supports summarizing features into higher-level groups. This enables taxonomic classification, for comparison purposes. The output of Woltka’s classification function and that of SHOGUN’s “assign_taxonomy” function are identical. Woltka supports three formats of classification systems: 1) the Greengenes-style lineage strings (supported by programs such as QIIME 2 (1), MetaPhlAn (21) and GTDB-tk (46)); 2) The NCBI-style taxonomy database (47) (a.k.a. “taxdump”, supported by programs such as Kraken 2 (48), Centrifuge (20) and DIAMOND (45)); 3) One or multiple plain mappings of child-to-parent classification units.

### Deployment

The Woltka program has been incorporated in the Qiita web analysis platform (https://qiita.ucsd.edu/) (13), as part of the standard operating procedure for analyzing shotgun metagenomic data (qp-woltka, code hosted at: https://github.com/qiita-spots/qp-woltka). It can be directly launched from the graphic user interface. A job array system is used to parallelize analyses on a per-sample base to maximize processing speed. Each process uses eight cores of an Intel E5-2640 v3 CPU and 90 GB DDR4 memory. Two reference genome databases are available for user choice: 1) The “Web of Life” (WoL) database (10), with 10,575 bacterial and archaeal genomes that were evenly sampled through an algorithm. 2) The reference and representative genomes of microbes defined in NCBI RefSeq release 200 (8). The subsequent community ecology analyses based on the OGU table are also available from Qiita. The WoL reference phylogeny is available for choice for phylogenetic analyses (such as UniFrac (4)).

This system allowed us to re-analyze all metagenomic datasets hosted on Qiita (totaling 143 studies and 57,063 samples, as of Mar 3, 2021) to generate OGU tables as well as tables at multiple taxonomic ranks, which are ready for subsequent meta-analysis by Qiita users. Although runtime varies by sample size, the average wall clock time for analyzing one metagenomic sample (including sequence alignment against WoL using Bowtie2 and feature table generation using Woltka) was 13.8 minutes in this large effort.

### The HMP dataset

The Human Microbiome Project (HMP) (17) dataset was downloaded from the official website (https://www.hmpdacc.org/hmp/). It contains 241 samples of 100 bp paired-end whole genome sequencing (WGS) reads. The sequencing data were already processed to remove human contamination and low-quality regions. We dropped samples with less than 1M paired-end reads, leaving 210 samples. They were randomly subsampled to 1M paired-end reads per sample. These samples represent both male (*n*=138) and female (*n*=72) human subjects. They represent seven body sites: stool (*n*=78), tongue dorsum (*n*=42), supragingival plaque (*n*=33), buccal mucosa (*n*=28), retroauricular crease (*n*=13), posterior fornix (*n*=10), and anterior nares (*n*=6).

### Taxonomic profiling

In comparison with the OGU method, we performed taxonomic profiling on the shotgun metagenomic data using four existing methods, specified as below. The default parameters were used for all programs. To maximize comparability, we used the WoL reference genome database (10) for all methods, except for MetaPhlAn (because it uses a special marker gene database which is difficult to customize).

1. SHOGUN: SHOGUN v1.0.8 (19), which calls Bowtie2 v2.4.1 to perform sequence alignment.
2. Bracken: Bracken v2.5 (18) on the results of Kraken v2.0.8 (48).
3. Centrifuge: Centrifuge v1.0.3 (20).
4. MetaPhlAn: MetaPhlAn v2.6.0 (21) with its database (mpa_v20_m200). Results (relative abundances) were normalized to counts per million sequences.

### Beta diversity analysis

Beta diversity analysis of the HMP dataset was performed using QIIME 2 (1), following recommended protocols (49). Specifically, beta diversity distance matrices were constructed using the “`qiime diversity beta`” command with Jaccard and Bray-Curtis metrics, and using the “`qiime diversity beta-phylogenetic`” command (50) with unweighted UniFrac and weighted UniFrac metrics, based on the WoL reference phylogeny. Principal coordinates analysis (PCoA) was performed using the “`qiime diversity pcoa`” command. The correlation between biological factors (body site and host sex) and beta diversity was assessed using the PERMANOVA test, through the command “`qiime diversity adonis`”, with 999 permutations (the default setting).

### Site clustering by environment

In the HMP study, we quantified the proximity of the three oral sites (tongue dorsum, supragingival plaque, and buccal mucosa) as compared with the four non-oral sites (stool, retroauricular crease, posterior fornix, and anterior nares) as follows: For each sample in the three oral sites, we calculated the beta diversity distance to all samples in all but the current site. We then separated these distances into oral (i.e., the two oral sites other than the current one) and non-oral (i.e., the four non-oral sites). We calculated the ratio of the mean distance of the former versus the latter. Finally we reported the distribution of the mean ratios of all oral samples.

### Phylogenetic factorization

We performed phylogenetic factorization as implemented in Phylofactor v0.0.1 to infer phylogenetic clades (“factors”) that are differentially abundant between male and female subjects. Two samples with less than 100,000 OGU counts were excluded from the analysis. OGUs with relative abundance below 0.01% were dropped from each sample, and OGUs present in fewer than two samples were also excluded. We built an explained variance-maximizing (the choice parameter was set to “var”) Phylofactor model using the OGU table and the WoL phylogeny. We specified the model to return 20 factors. They were labeled by the taxonomic annotation of the corresponding phylogenetic clades as provided in the WoL database. The results were visualized with EMPress. In each factor, we tested the differences in male vs female subjects by comparing the ILR-transformed vectors corresponding to each sample group using a two-tailed independent samples *t*-test.

### Subsampling of OGU tables

To assess the impact of sampling depth on analysis results, we randomly subsampled the OGU tables to lower depths (sum of OGU frequencies per sample). This process mimicked lower sequencing depths in the original data, because the sum of OGU frequencies is nearly equal to the number of aligned sequences (see above). This process further considered the unaligned part of the sequencing data. For example, if *m* out of *n* sequences in a sample were aligned to at least one reference genome (therefore the sum of OGU frequencies was *m*), we added an extra “unaligned” feature of frequency of *n* - *m* to the OGU table, prior to random subsampling, and removed this feature after sampling.

### The FINRISK 2002 datasets

The FINRISK 2002 is a large, well-phenotyped, and representative cohort based on a stratified random sample of the population aged 25 to 74 years from specific geographical areas of Finland (27). All volunteer participants took a self-administered questionnaire, physical measurements and collection of blood and stool samples. The microbiome data and metadata that support the findings of this study are available from the THL Biobank based on a written application and following relevant Finnish legislation. Details of the application process are described in the website of the Biobank: https://thl.fi/en/web/thl-biobank/for-researchers.

Paired 16S rRNA gene amplicon sequencing data and shotgun metagenomic sequencing data are available for 6,430 stool samples. The 16S rRNA data were demultiplexed, quality filtered, and denoised with deblur v1.1.0 (51), resulting in an average ASV frequency of 8,787 per sample, followed by normalization to 10,000 per sample. Taxonomic classification was performed using a pre-trained Naive Bayes classifier against the Greengenes 13_8 database at an OTU clustering level of 99%. Feature tables were rarefied to a sampling depth of 10,000. The shotgun metagenomic data were trimmed and quality filtered using Atropos v1.1.25 (52), resulting in an average of 1.07 million paired-end sequences per sample. They were aligned to the WoL database using SHOGUN v1.0.8. An OGU table was generated using the current approach. As a comparison, Bracken v2.5 with Kraken v2.0.8 were used to infer taxonomic profiles using the same WoL database. These analyses were the same as the corresponding analyses of the HMP shotgun metagenomic dataset, as described above.

### Supervised regression for age prediction

We performed machine learning analysis of microbial profiles derived from both 16S amplicon sequencing and shotgun metagenomics sequencing, at distinct levels of resolution. These included taxonomic ranks (phylum, class, order, family, genus and species) for both 16S rRNA and shotgun metagenomic data (the latter of which were inferred by either SHOGUN or Bracken), ASV for 16S rRNA data, and OGU for shotgun metagenomic data (inferred by SHOGUN with Woltka). In each profile, features with a study-wide prevalence less than 0.001 were excluded. Random Forest regressors for predicting chronological age were trained based on each profile with tuned hyperparameters with a stratified 5-fold cross-validation approach using R package ranger v0.12.1 (53). Each dataset was split into five groups with similar age distributions, and we trained the classifier on 80% of the data, and made predictions on the remaining 20% of the data in each fold iteration. We next evaluated the performance of age prediction using mean absolute error (MAE), which calculated as 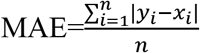, where *y* denotes the predicted age, *x* denotes the chronological age, and *n* is the total number of samples. Based on the MAE evaluation, we next determined the most predictive taxonomic levels derived from both 16S and shotgun metagenomics.

To identify the most important taxonomic features that contributed to the age prediction, we visualized the top-128 ranked important features by built-in Random Forest importance scores and their phylogenetic relationships using EMPress (54). We next performed the feature selection analysis to identify a set of important microbial features that can maximize the model performance. We built age regressors using a series of reduced sets (*n* = 2, 4, 8, 16, 32, 64, 128, 256, 512, 1024, and the number of all features) of the most predictive taxonomic features (namely, OGU) and compared their performance. The rationale is to observe the trough in MAE when additional features are added into the regression model.

### Statistics statement

All data analysis was performed using QIIME 2 release 2020.6. PERMANOVA was performed using the “adonis” command (which wraps the “adonis” function in vegan v2.5-6). Paired *t*-test was performed using the “ttest_rel” function in SciPy v1.4.1.

## Acknowledgements

We are grateful to Gabriel Al-Ghalith, Zachary Burcham, Jeff DeReus, Marcus Fedarko, Shalisa Hansen, Stefan Janssen, Emily Kobayashi, Evguenia Kopylova, Tomasz Kosciolek, Holly Lutz, Cameron Martino, Siavash Mirarab, James Morton, Oriane Moyne, Wayne Pfeiffer, Daniel Roush and Se Jin Song for valuable testing of the methodology, insightful discussions on this study and additional assistance.

This work is supported in part by an Arizona State University start-up grant (to Q.Z.), Sloan Foundation G-2017-9838, IBM Artificial Intelligence for Healthy Living A1770534, DARPA JUMP/CRISP, NIH P30DK120515, DP1AT010885, U19AG063744, U24CA248454, Emerald Foundation Distinguished Investigator Award, Crohn’s and Colitis Foundation 675191, NSF RAPID 2038509, IBM Research AI through the AI Horizons Network and the UC San Diego Center for Microbiome Innovation (to S.H., I.M., Y.V.-B., and R.K.). G.D.S.-P. is supported by a fellowship from the National Institutes of Health (F30 CA243480). T.N. was funded by the Emil Aaltonen Foundation, the Finnish Medical Foundation, the Finnish Foundation for Cardiovascular Disease, and the Academy of Finland (grant 321351). V.S. was supported by the Finnish Foundation for Cardiovascular Research. This work used the Comet supercomputer at the San Diego Supercomputer Center through allocation BIO150043 through the Extreme Science and Engineering Discovery Environment (XSEDE).

Q.Z. and R.K. conceived the project. Q.Z. led the development of the methodology and software. S.H. and Q.Z. led the analysis and interpretation of the datasets presented in this article. S.H., A.G., D.M. and Y.V.-B. contributed to the design of the method. A.G., D.M. and G.A. contributed to the development of the software. G.D.S.-P., A.D.S., P.D., F.L. contributed to the test of the method. P.B.-F., A.S.H., G.M., T.N., L.L., V.S. and M.J. contributed to data curation. A.G., I.M., J.Y., Y.V-B. and J.K. contributed to data analysis. N.H., G.D.S.-P., A.S.H., G.M., T.N., L.L., V.S., H.-C.K., M.J., M.I., J.A.G. and R.K. contributed to result interpretation. R.K. and Q.Z. managed the project. All the authors contributed to the composition and discussion of the manuscript.

We declare that we have no competing interests.

